# Proteomics Using Protease Alternatives to Trypsin Benefits from Sequential Digestion with Trypsin

**DOI:** 10.1101/2020.01.30.927046

**Authors:** Therese Dau, Giulia Bartolomucci, Juri Rappsilber

## Abstract

Trypsin is the most used enzyme in proteomics. Nevertheless, proteases with complementary cleavage specificity have been applied in special circumstances. In this work, we analyzed the characteristics of five protease alternatives to trypsin for protein identification and sequence coverage when applied to *S. pombe* whole cell lysates. The specificity of the protease heavily impacted on the number of proteins identified. Proteases with higher specificity let to the identification of more proteins than proteases with lower specificity. However, AspN, GluC, chymotrypsin and proteinase K largely benefited from being paired with trypsin in sequential digestion, as had been shown by us for elastase before. In the most extreme case, pre-digesting with trypsin improves the number of identified proteins for proteinase K by 731 %. Trypsin pre-digestion also improved the protein identifications of other proteases, AspN (+62 %), GluC (+80 %) and chymotrypsin (+21 %). Interestingly, the sequential digest with trypsin and AspN yielded even higher number of protein identifications than digesting with trypsin alone.

## INTRODUCTION

Trypsin is the protease of choice for mass spectrometry (MS)-based proteomics. It cleaves carboxyterminal of Arg and Lys residues, resulting in a positive charge at the peptide C-terminus, which is advantageous for MS analysis ^1,2^. Nevertheless, other proteases are frequently used to obtain complementary data ^3,4^.

Among these, AspN and GluC target acidic amino acid residues (**Figure 1a**). Both enzymes generate peptide mixtures of comparable complexity to that of trypsin and have been successfully used in many studies ^4–7^. Chymotrypsin, which targets primarily aromatic residues, has also been used ^7–9^. In contrast, broad specificity proteases are much less widely used in proteomics. This is likely due to the high complexity of the peptide mixtures that they generate. To our knowledge their application has been limited to pre-fractionated samples. Proteinase K for example, was used, to “shave” surface exposed loops from proteins in membrane vesicles ^10,11^

**Figure 1:**
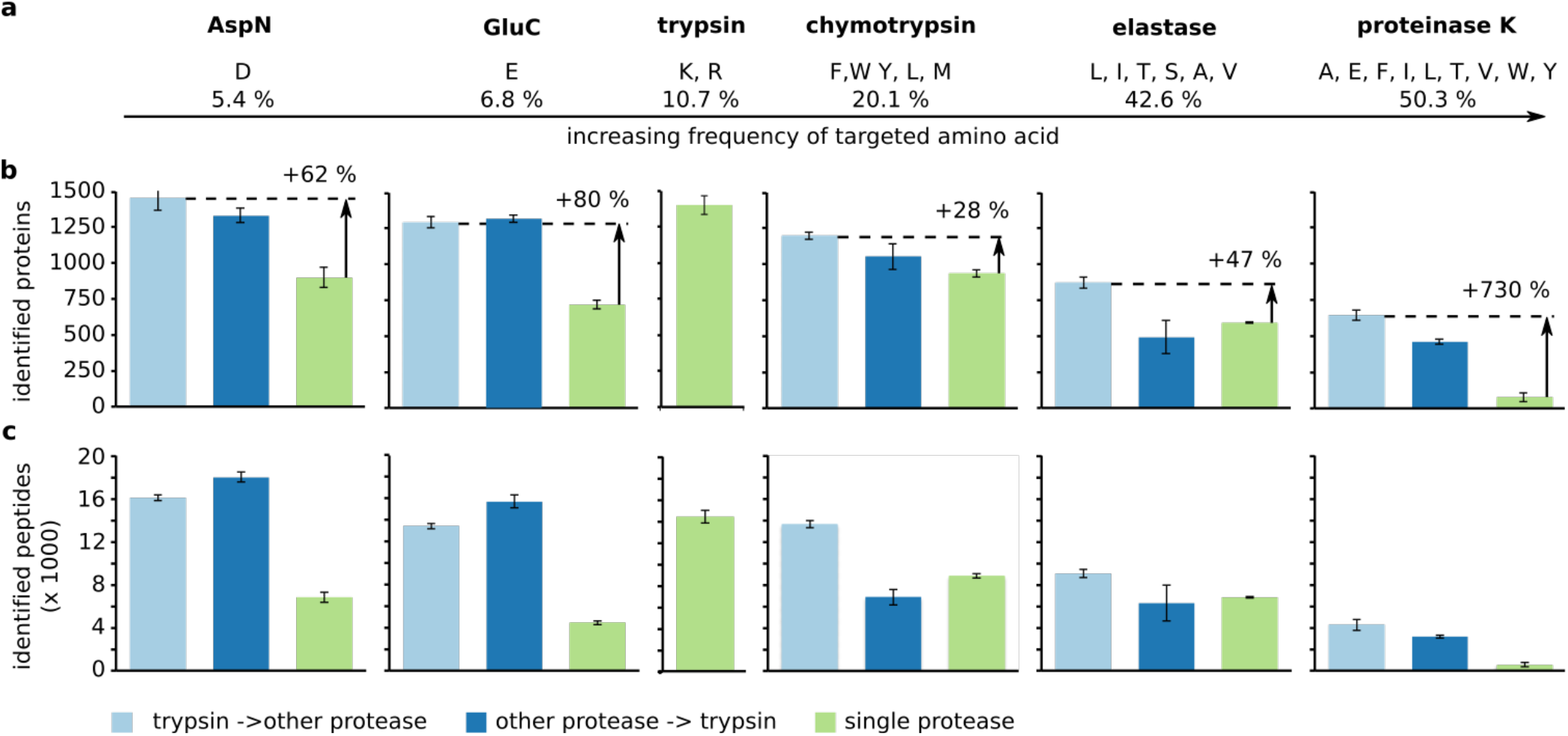
Impact of different proteases and protease combinations on the identification of proteins and peptides. (a) Frequency of the amino acids targeted by AspN, GluC, chymotrypsin, elastase and proteinase K according to the UniProtKB/TrEMBL release report. Number of (a) proteins and (b) peptides identified with different protease combinations. Trypsin-other protease, light blue; other protease-trypsin, dark blue; single protease, green. Error bars are standard deviation (SD) of at least 5 independent digestion experiments.

Our group has previously shown that the number of identified peptides, when using alternatives to trypsin, could largely be improved by a sequential combination with trypsin. This includes AspN, GluC, chymotrypsin and elastase for the detection of crosslink sites ^12–14^ and elastase applied to *S. pombe* whole cell lysates ^14^. The sequential digestion increased the number of identified crosslinks up to 19-fold for the Taf4-12 complex compared to digesting with elastase alone ^14^. Introducing positively charged C-termini through trypsin improves the detection of previously non-tryptic peptides. Importantly, smaller peptides are protected from the second protease ^12,14^. Thus, use of two proteases does not lead to the very small peptides that *in silico* digestion would predict. In consequence, using elastase after trypsin does not lead to the same peptide complexity as using elastase alone.

In this study, we analyzed whether the introduction of trypsin in a sequential digest, might improve the application of AspN, GluC, chymotrypsin, and proteinase K on unfractionated *S. pombe* lysate.

## METHODS

### Public datasets

The data on trypsin, elastase, trypsin-elastase and elastase-trypsin were taken from our previous work ^14^ and retrieved from PRIDE with the data set identifier PXD011459.

### Sample preparation

One gram of frozen and ground *S. pombe* cells were resuspended in 2 ml RIPA (Sigma-Aldrich, St. Louis, MO) supplemented with cOmplete (Roche, Basel). To remove the cell debris the samples were centrifugated at 1200 g for 15 minutes. The lysates were subjected to gel electrophoresis on a 4-12 % Bis-Tris gel (Life Technologies, Carlsbad, CA) for 5 minutes and stained using Imperial Protein Stain (Thermo Fisher Scientific, Rockford, IL). After excising the stained gel area as a single fraction, the proteins were first reduced with dithiothreitol and then alkylated with iodoacetamide.

The first protease (trypsin (13 ng/μl), elastase (15 ng/μl), AspN (1:100), GluC (1:50), chymotrypsin (1:50) and proteinase K (1:50) were incubated for 16 hours at 37°C (besides chymotrypsin at RT). The second protease was added for 4 hours at 37°C (besides elastase for 30 minutes).

We used a standardized protocol to desalt and concentrate the peptides on C18 StageTips for subsequent analysis ^15,16^.

### LC-MS/MS

All samples were analyzed on a linear iontrap-orbitrap mass spectrometer (Orbitrap Elite, Thermo Fisher Scientific, Rockford, IL) coupled online to a liquid chromatograph (Ultimate 3000 RSLCnano Systems, Dionex, Thermo Fisher Scientific, UK) with a C18-column (EASY-Spray LC Column, Thermo Fisher Scientific, Rockford, IL). The flow rate was 0.2 μL/min using 98 % mobile phase A (0.1 % formic acid) and 2 % mobile phase B (80 % acetonitrile in 0.1 % formic acid). To elute the peptides, the percentage of mobile phase B was first increased to 40 % over a time course of 110 minutes followed by a linear increase to 95 % in 11 minutes. Full MS scans were recorded in the orbitrap at 120,000 resolution for MS1 with a scan range of 300-1700 m/z. The 20 most intense ions (precursor charge ≥2) were selected for fragmentation by collision-induced disassociation and MS2 spectra were recorded in the ion trap (20,000 ions as a minimal required signal, 35 normalized collision energy, dynamic exclusion for 40 s).

### Data analysis

MaxQuant software ^17^ (version 1.5.2.8) employing the Andromeda search engine ^18^ in combination with the PombeBase database ^19^ was used to analyze the samples. The following parameters were used for the search: carbamidomethylation of cysteine as a fixed modification, oxidation of methionine as a variable modification, MS accuracy of 4.5 ppm and MS/MS tolerance of 20 ppm. Up to 6 miscleavages were allowed for digests involving trypsin, AspN, GluC or chymotrypsin and up to 10 miscleavages for digests containing elastase or proteinase K. Frequency of amino acids were taken from the statistics of the UniProtKB/TrEMBL protein database release 2019_11 (https://www.ebi.ac.uk/uniprot/TrEMBLstats).

## RESULTS AND DISCUSSION

Lysate from *S. pombe* was digested either with trypsin, AspN, GluC, chymotrypsin, elastase or proteinase K. We also combined each of the proteases other than trypsin in a sequential digest with trypsin as either the first or second protease.

Adding trypsin to the digest with AspN and GluC improved the protein (AspN = 899 ± 69, trypsin-AspN = 1455 ± 85, AspN-trypsin = 1331 ± 50, GluC = 719 ± 28, trypsin-GluC = 1294 ± 37, GluC-trypsin = 1319 ± 25) identification (**Figure 1b**). Peptide identifications also improved (AspN = 6828 ± 514, trypsin-AspN = 16087 ± 327, AspN-trypsin = 17968 ± 470, GluC = 4467 ± 182, trypsin-GluC = 13461 ± 260, GluC-trypsin = 15713 ± 600) (**Figure 1c**). The order of proteases had only a minor influence on the identifications.

Using trypsin prior to chymotrypsin or elastase also improved the identification of proteins (chymotrypsin = 938 ± 27, trypsin-chymotrypsin = 1200 ± 25, elastase = 593 ± 7, trypsin-elastase = 874 ± 40) and peptides (chymotrypsin = 8818 ± 232, trypsin-chymotrypsin = 13611 ± 346, elastase = 6821 ± 84, trypsin-elastase = 9039 ± 374). Using trypsin as the second protease had only a minimal effect on the protein (chymotrypsin-trypsin = 1056 ± 91, elastase-trypsin = 492 ± 115) and peptide identification (chymotrypsin-trypsin = 6869 ± 744, elastase-trypsin = 6280 ± 1680).

Interestingly, digesting with trypsin alone did not give the highest number of protein (1403 ± 65) and peptide (14410 ± 571) identifications. We identified more proteins (+4 %) and peptides (+12 %), when trypsin was followed by AspN.

The biggest impact of sequential digestion with trypsin was seen on the performance of proteinase K. Using proteinase K alone led to very few identifications of proteins (proteinase K = 78 ± 33) and peptides (proteinase K = 527 ±179). This might be due to very short peptides being generated by proteinase K, which cleaves carboxyterminal of half of all the amino acids. Alternatively, or in addition, the high complexity of the peptide mixture generated by proteinase K might reduce identification rates. Surprisingly, adding trypsin to the proteinase K digest increased the number of identifications for proteins (proteinase K-trypsin = 461 ± 17) and peptides (proteinase K-trypsin = 3169 ± 194). Using trypsin prior to proteinase K further improved on these results as this led to the identification of 8x more proteins (646 ± 36) and 8x more peptides (4279 ± 530) compared to proteinase K alone.

In summary, AspN, GluC and proteinase K profited most of the five tested proteases from the addition of trypsin. The underlaying reasons for the observed gains are likely different. AspN and GluC have a low amount of available cleavage sites and therefore generate relatively long peptides. Many of these will be unfavorably long for mass spectrometric detection. In addition, they are missing a terminal positive charge. Adding trypsin introduces such a C-terminal charge and shortens very long peptides, both enhancing peptide detection in MS analysis.

While AspN and GluC are very specific proteases, over 50 % of the residues are potential cleavage sites for proteinase K. The problem for proteinase K is therefore not a lack of cleavage sites. Adding trypsin to proteinase K increased identifications and thus ruled out the possibility that peptides generated by proteinase K alone, at least under standard conditions, are generally too short for proteomics. If therefore complexity of a proteinase K digest is the reason for the low identification yields of proteinase K, then the addition of trypsin must reduce this complexity. Adding trypsin might unify “ragged” proteinase K peptides that share either the N- or C-terminus but have different length (**Figure 2a**). In this way trypsin leads to a concentration increase of peptides by reducing sample complexity. At least when trypsin is used first an additional mechanism must be considered that was previously described for sequential digestion ^12,14^. The second enzyme does not cleave shorter peptides with high efficiency effectively leading to short tryptic peptides being protected from proteinase K. In either case, the complexity that is normally introduced through proteinase K is reduced by the tryptic treatment.

**Figure 2:**
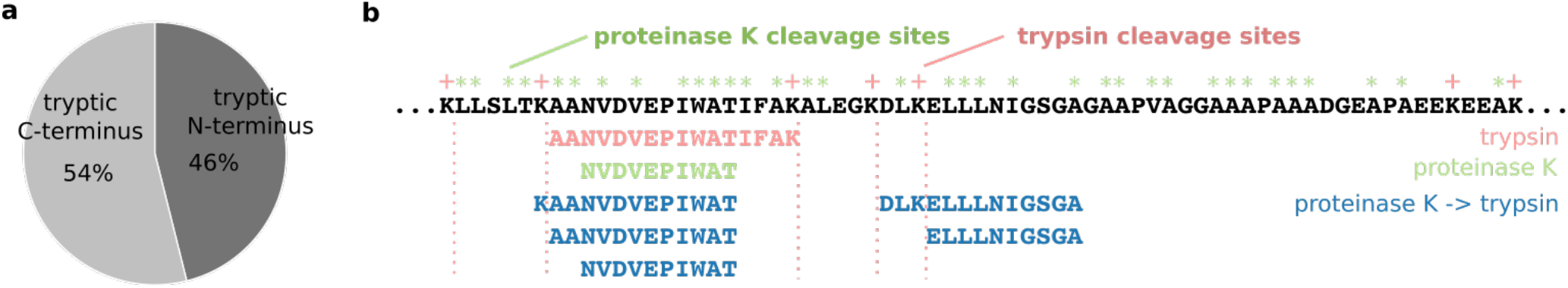
(a) Comparison of semi-tryptic peptides with a tryptic N or C-terminus after digesting whole *S. pombe* with proteinase K followed by trypsin. (b) Peptides that have been identified in the N-terminal region of 60S acidic ribosomal protein P1-alpha 1 (26-94) with either trypsin, proteinase K or proteinase K followed by trypsin. Trypsin, red; proteinase K, green; proteinase K-trypsin, dark blue.

**Figure 3:**
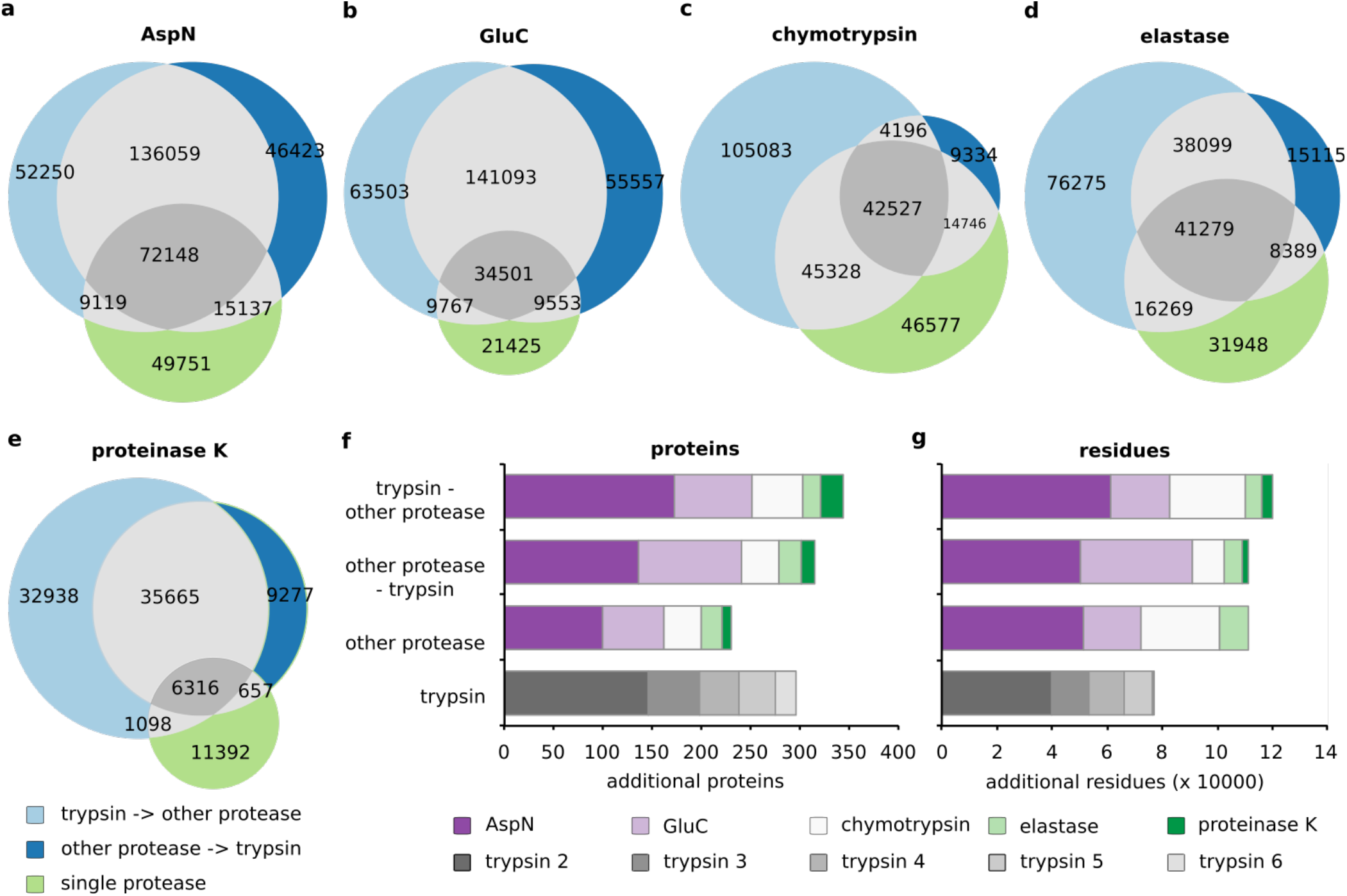
Comparison of residues covered with the three protease combinations for (a) AspN, (b) GluC, (c) chymotrypsin, (d) elastase, (e) proteinase K. The result of all five experiments of each condition was used for the analysis. Trypsin-other protease, light blue; other protease-trypsin, dark blue; single protease, green. Comparison of (f) proteins and (g) residues gained on top of a tryptic digest trough sequential digestion variants, parallel digestion and replica of tryptic digestion. AspN, dark violet; GluC, light violet; chymotrypsin, white; elastase, light green; proteinase K dark green.

End trimming and short peptide protection alone are likely not the sole explanation. We observed previously that among all observed mixed-protease action peptides, i.e. those peptides that were generated by trypsin action on one end and another protease at the other end, there is a misbalance: tryptic C-termini are more prevalent than N-termini generated by trypsin (semi-tryptic peptides with tryptic N-terminus = 652 ± 42, semi-tryptic peptides with tryptic C-terminus = 763 ± 15) (**Figure 2a**). This means, also the improved observability of peptides with a basic C-terminal residue contributes to the observed effect of sequential digestion on identification rates.

As an example, we analyzed the 60S acidic ribosomal protein P1-alpha 1 (**Figure 2b**). There are no trypsin cleavages sites between residues 56 to 90, so this region is not covered when trypsin is used alone. Digesting with proteinase K alone did not improve the coverage for this region, although or possibly because every other residue is a potential cleavage site for proteinase K. Peptides from this region could only be identified, when proteinase K and trypsin were used in a sequential digest.

We then wondered how far the proteins and peptides that were observed in the different uses of proteases alone or in combination with trypsin, covered different sequence space. We measured this in number of unique residues. As one would expect, this followed the same trends seen for protein and peptide identifications. For AspN and GluC the largest number of residues was covered, when trypsin was used following the other protease (**Figure 2a, b**). For chymotrypsin, elastase and proteinase K the inverse order, i.e. trypsin first yielded the larger coverage (**Figure 2c, d, e**). Nonetheless, the different conditions yielded substantial non-overlap. When combining the results of two digestion conditions one would combine the data obtained by the protease alone with that of a trypsin-first sequential digest. Their overlap is substantially smaller (4 ± 2 % to 38 ± 1 %) than what we observed here for trypsin replicas (83 ± 2 %).

Finally, we analyzed the gain in identified proteins and residues when using different combinations of digestion conditions (**Figure 2f, g**). We combined the results of either five replicas of trypsin, trypsin followed by either of the five other proteases or either of the five other proteases followed by trypsin. An initial trypsin digest served as reference, in which 1484 proteins and 202,556 residues were identified. This followed the rationale that one would always use trypsin for an initial analysis, although trypsin followed by AspN in a sequential digest consistently gave here higher protein and peptide identifications. The highest number of complementary proteins (344) and residues (119,763) were identified when trypsin was used first in a sequential digest in combination with either of the five other proteases, followed by the inverted set-up in which trypsin was used last (proteins = 315, residues = 111,126). Digestions with non-tryptic proteases alone were outperformed by trypsin replicas in terms of protein identification (230 versus 296) but not in terms of residue coverage (111,069 versus 76,645).

## CONCLUSION

In this study, we investigated the impact of adding trypsin to other proteases in proteomics. Sequential digestion has been used before ^5,6,20^ and we here add a systematic evaluation of different protease combinations. Protein and peptide identifications improved when combining any of the tested proteases with trypsin. This is in line with previous studies on cross-linking identification, which benefited from the sequential digest with trypsin ^12,14^. In the most extreme case, the sequential digest with trypsin and AspN outperformed results obtained by trypsin alone. This effect is relatively small and due to cost considerations, trypsin will remain the protease of first choice in proteomics also after our study. However, situations where alternative proteases are currently used could in future benefit from adding a sequential digestion step with trypsin. As trypsin is compatible with the buffer conditions of the tested proteases this requires no other additional step than adding trypsin.

## AUTHOR INFORMATION

### Author Contributions

T.D. and J.R. conceived this study and interpreted data, T.D and G.B. conducted all experiments. T.D. and J.R. wrote the manuscript with input from G.B. All authors have given approval to the final version of the manuscript.

### Accession Codes

The mass spectrometry proteomics data have been deposited to the ProteomeXchange Consortium ^21^ via the PRIDE partner repository with the data set identifier PXD011459 ^14^ and PXD017321.

### Notes

The authors declare no competing interests.

## ACKNOWLEDGMENT

This work was supported by a research stipend to T.D. (DA 1861/2-1) from the Deutsche Forschungsgemeinschaft and by the Wellcome Trust through a Senior Research Fellowship to J.R. (103139) and a multiuser equipment grant (108504). The Wellcome Centre for Cell Biology is supported by core funding from the Wellcome Trust (203149).

---

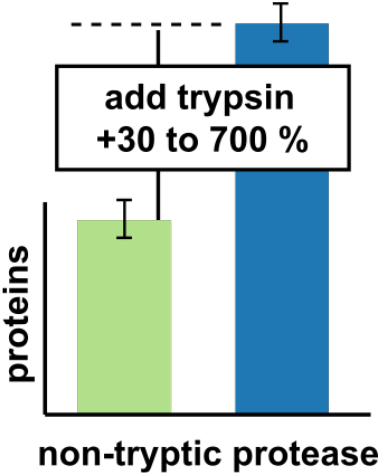

